# 12S Gene Metabarcoding with DNA Standard Quantifies Marine Bony Fish Environmental DNA, Identifies Threshold for Reproducible Amplification, and Overcomes Distortion Due to Non-Fish Vertebrate DNA

**DOI:** 10.1101/2022.07.29.502053

**Authors:** Mark Y. Stoeckle, Jesse H. Ausubel, Michael Coogan

**Author notes:** AUTHOR CONTRIBUTIONS STATEMENT Designed study: MYS Updated bioinformatics: MC Collected samples: MYS, JHA Analyzed samples: MYS Analyzed data: MYS, JHA Prepared figures: MYS Wrote first draft of manuscript: MYS Revised manuscript: MYS, JHA, MC.

## Abstract

Single-species PCR assays accurately measure eDNA concentration. Here we test whether multi-species PCR, i.e., metabarcoding, with an internal standard can quantify eDNA of marine bony fish. Replicate amplifications with Riaz 12S gene primers were spiked with known amounts of a non-fish vertebrate DNA standard, indexed separately, and sequenced on an Illumina MiSeq. Fish eDNA copies were calculated by comparing fish and standard reads. Relative reads were directly proportional to relative DNA copies, with average and maximum variance between replicates of about 1.3- and 2.0-fold, respectively. There was an apparent threshold for consistent amplification of about 10 eDNA copies per PCR reaction. The internal DNA standard corrected for distortion of read counts due to non-fish vertebrate DNA. To assess potential amplification bias among species, we compared reads obtained with Riaz 12S primers to those with modified MiFish primers. Our results provide evidence that Riaz 12S gene metabarcoding with an internal DNA standard quantifies marine bony fish eDNA over a range of about 10 to 5,000 copies per reaction, without indication of significant PCR bias among teleost species. In mid-Atlantic coastal samples, eDNA rarity was the main limitation to reproducible detection and quantification, and this was partly overcome by increasing the amount of a DNA sample amplified. Our findings support incorporating a DNA standard in 12S metabarcoding to help quantify eDNA abundance for marine bony fish species.

## INTRODUCTION

Addressing human impacts on the ocean calls for regular monitoring of marine life (Halpern et al., 2008; Lubchenko, Haugan, & Pangestu, 2020). Accurate assessment of marine fish populations, for example, enables effective fisheries management, helps gauge value of Marine Protected Areas (MPAs), and helps reveal ecological footprints of maritime industries including aquaculture and wind farms (Ahmadia et al., 2015; Fernandez et al., 2001; Hollingworth, 2000). Traditional survey methods for monitoring marine fish—capture, sonar, and visual—are relatively expensive and need specialized equipment and trained personnel. In some settings, traditional surveys may be harmful or otherwise unsuitable. For instance, bottom trawls may damage sea floor habitat and cannot be deployed at rocky sites (de Groot, 1984). Environmental DNA offers an informative addition to current survey techniques. Advantages include relatively low cost, modest field equipment, performance by a wide variety of personnel, harmlessness to environment, and applicability in difficult environments (Bourlat et al., 2013; Hansen, Bekkevold, Clausen, & Nielsen, 2018). Inherent disadvantages are absent information on organism size, life stage, sex, and health. Despite limitations, eDNA appears poised to become a routine monitoring tool for marine life (Gilbey et al., 2021; Hinz, Coston-Guarini, Marnane, & Guarini, 2022). Current challenges to wider adoption include lack of standardized methods, incomplete DNA reference libraries, and, most importantly, limited understanding of how eDNA abundance relates to organism abundance under various environmental conditions (Allan, Zhang, Lavery, & Govindarajan, 2020; Andruskiewiz et al., 2019; Collins et al., 2018; Jeunen et al., 2019).

Advancing knowledge of how fish species eDNA abundance informs on fish species abundance has two essential components. The first is accurately measuring the concentration of a species eDNA in collected water samples. The second is analyzing how that concentration relates to the species local abundance. This latter component can be challenging regarding marine fish, as there is no gold standard for abundance—all survey methods have biases (Arregúin-Sánchez, 1996; Frazier, Greenstreet, & Piet, 2007). In this report we focus on the first step, accurate measurement, and test whether 12S gene metabarcoding incorporating a DNA standard can be a quantitative tool for marine bony fish eDNA. Metabarcoding uses broad-range primers that amplify the target gene of multiple species in a taxonomic group, such as vertebrates, and applies Next Generation Sequencing (NGS) to sequence the resulting mixture of amplified DNA (Taberlet, Bonin, Zinger, & Coissac, 2018). Bioinformatic processing bins NGS output, called reads, into amplicon sequence variants (ASVs), and identifies ASVs by matching to a genetic reference library (Callahan et al., 2016). Metabarcoding is often considered a qualitative tool best suited for presence/absence surveys (Bush et al., 2020; Leray & Knowlton, 2015; Turunen et al., 2021). Hindrances to quantitative metabarcoding include differing PCR bias among species, distortion of read counts due to amplification of non-target DNA, and the fact that metabarcoding PCR tends to generate the same number of reads unrelated to the amount of eDNA added (Kelly, Shelton, & Gallego, 2019; Krehenwinkel et al., 2017). In contrast, qPCR and other single-species techniques provide reliable measures of eDNA concentration (Andruszkiewicz, Yamahara, Closek, & Boehm, 2020; Doi et al., 2015; Shelton et al., 2022a). However, it may be clumsy to develop and apply single species assays covering the multitude of taxa that are actively managed or otherwise of interest. For example, in our coastal mid-Atlantic study area, about 30 fish species are subject to catch quotas or are monitored due to threatened status (Atlantic States Marine Fisheries Commission, 2022). For more general ecosystem assessment such as for MPAs, taxonomically broad-range monitoring with metabarcoding would be helpful. Regional targets might include the approximately 100 fish species captured annually in bottom trawl surveys, or the over 400 species recorded in coastal waters (Able, 1992).

In this report we analyze bony fish eDNA in water samples collected during New Jersey Ocean Trawl Survey (NJOTS) and from an ongoing environmental survey of shoreline sites in coastal New Jersey, New York, and Massachusetts (Hinks & Berry, 2020; Stoeckle et al., 2021,2022). We test whether adding a DNA standard to metabarcoding PCRs enables reproducible measurement of marine bony fish eDNA concentration. Advantages of our regional focus include long-term data on fish abundance obtained with NJOTS (Levesque, 2019), an excellent DNA reference library for commonly encountered fish species, and, among bony fish, highly conserved binding sites for mitochondrial 12S gene metabarcoding primer sites. Marine fish differ widely in abundance. For example, in NJOTS, monthly biomass per species ranges over four or five orders of magnitude (e.g., Fig. 1). With this in mind, we aimed for eDNA measurement accurate within one order of magnitude. Ultimately, of course, greater precision is desirable, but even order-of-magnitude assessments could be useful for ecosystem monitoring, for example. To better understand sources of variation and the value of replicate PCR, all replicates were analyzed individually. As part of assessing metabarcoding quantification, we ask three related questions: First, is there a lower limit to reproducible detection? Second, does DNA standard overcome distortion due to non-fish vertebrate DNA? Third, how important is PCR bias in quantifying marine bony fish eDNA?

**Fig. 1.**
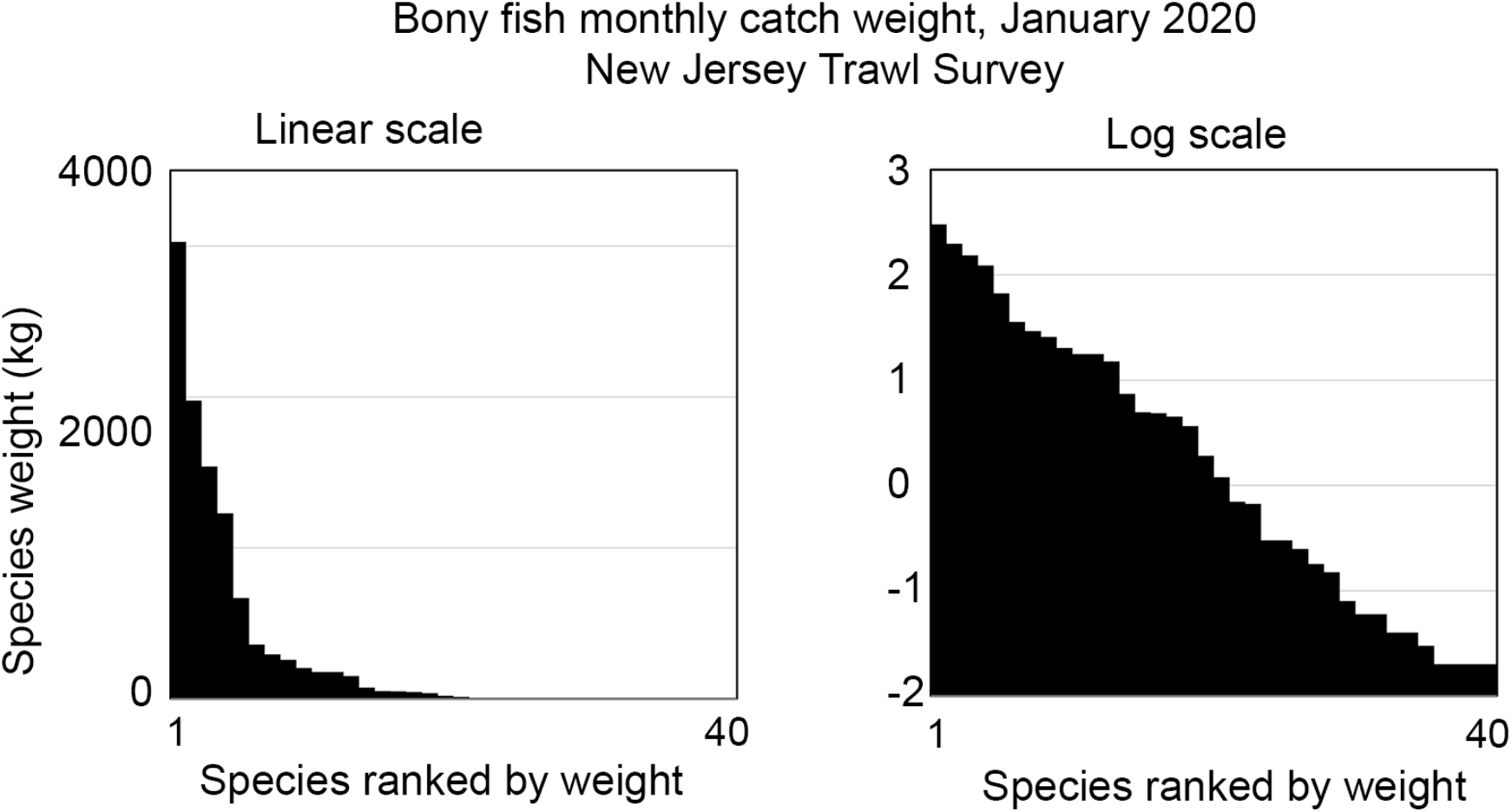
Mid-Atlantic bony fish abundance differs over multiple orders of magnitude. Source data are in Supporting Information Table 1.

## MATERIALS AND METHODS

Most of the methods employed in this study have previously been described in detail (Stoeckle et al., 2021, 2022). These are briefly summarized here, and additional details are in Supporting Information Table 2.

### Water collection, filtration, DNA extraction

Standard water collection volume was one liter. The data presented were obtained from 19 NJOTS and 23 environmental survey samples, plus tap water controls. Filtration was done using a 47 mm diameter, 0.45 μM pore size nitrocellulose filter, and filters were stored at −20°C. DNA was extracted with DNeasy PowerSoil Pro Kit (Qiagen), recovered in 100 μl Buffer C6, and stored at −20°C. Negative filtration and extraction controls were prepared from one liter samples of laboratory tap water and were processed using the same equipment and procedures as for field samples. No animals were housed or experimented upon as part of this study. No endangered or protected species were collected.

### PCR

Metabarcoding PCR reactions were carried out in 25 μl total volume with TaKaRa High Yield PCR EcoDry™ Premix. Standard conditions were 5 μl of extracted DNA or 5 μl of molecular biology grade water, and 200 nM Illumina-tailed Riaz 12S primers (IDT) (Fig. 2) (Riaz et al., 2011). Where noted, modified MiFish-U-F/R2 primers were used (Miya et al., 2015; Stoeckle et al., 2022). The modified set has reduced off-target amplification of bacterial 16S gene, enabling amplicon sequencing without gel purification. Primer sequences and thermal cycling conditions are shown in Supporting Information Table 2. Negative control reactions were included in all amplification sets. 5 μl of each reaction mix were run on a 2.5% agarose gel to assess amplification, and the remaining 20 μl were diluted 1:20 in Buffer EB (Qiagen) to be used as template for indexing. Indexing was done with Nextera XT kit and Cytiva PuReTaq Ready-To-Go PCR beads, using 5 μl of diluted primary PCR product as template (thermal cycling protocol in Supporting Information Table 2).

**Fig. 2.**
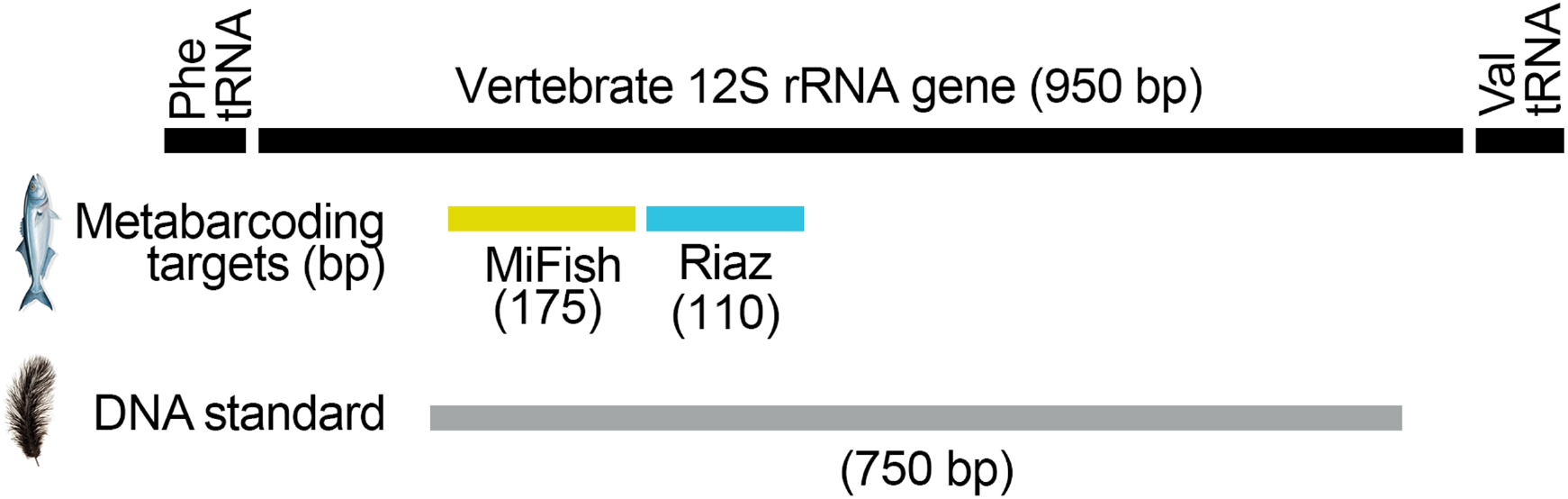
Targeted gene regions. Schematic of vertebrate 12S gene with metabarcoding targets and segment used for standard shown.

### Next-generation sequencing, bioinformatic analysis

Sequencing was performed at GENEWIZ on an Illumina MiSeq, 2 x 150 bp (2 x 250 bp for MiFish libraries), with 10% PhiX added to each run. Bioinformatic analysis was performed on Illumina FASTQ files using a DADA2 pipeline (Callahan et al., 2016; Callahan, McMurdie, & Holmes, 2017). Taxon assignments were generated by comparison to an internal 12S gene reference library for regional fishes and other commonly amplified vertebrate ASVs (Supporting Information File 1). In addition, all ASVs were manually submitted to GenBank to recover overlooked matches to 12S gene sequences not included in internal library. All identifications were based on 100% match to a reference sequence. Some near shore samples generated matches to non-local fish species, consistent with wastewater origin (Fujii et al., 2019). These were excluded from analysis. In addition, given that cartilaginous fish eDNA amplifies poorly with both primer sets used in this study, elasmobranch detections were set aside. Tap water eDNA and reagent grade water libraries were negative for fish ASVs after filtering DADA2 output tables as previously described. Filtering consisted of excluding detections comprising less than 1/1000^th^ of the total for that taxon among all libraries in the run. For MiFish amplifications, all ASVs were manually submitted to GenBank. The data presented were obtained from 144 libraries generated from the 42 field samples plus controls, which were analyzed in five MiSeq runs together with other samples not reported here. The MiFish 12S gene segment distinguishes some species which have identical Riaz 12S gene sequences. To enable comparison of Riaz and MiFish primers, ASV tables were harmonized by combining MiFish reads for species with identical Riaz sequences (relevant species listed in Supporting Information Table 3).

### 12S gene DNA standards

We selected ostrich (*Struthio camelus*) and emu (*Dromaius novaehollandiae*) as internal standards, as DNA could be obtained from commercial food products (ostrich, American Ostrich Farms; emu, Newport Jerky Company), both were unlikely to be present in regional environmental samples, and Riaz 12S gene metabarcoding primer sites were identical to those in bony fish. Standards were generated with M13 tailed primers that amplify a 689 bp segment of 12S gene covering the Riaz primer target sites and flanking regions (Fig. 2). Primer sequences and thermal cycling parameters are in Supporting Information Table 2. Sanger sequencing confirmed 100% match to *S. camelus* or *D. novaehollandiae* reference sequences. PCR products were purified with AMPure beads at 1:1, concentration measured by Qubit, and a series of 10-fold dilutions were prepared in Buffer EB. Amounts added to replicates in this study ranged from 500 ag to 0.5 ag, corresponding to 600 to 0.6 copies, respectively. Linear regression and Fisher’s exact test analyses were made with Prism 8.

## RESULTS

To test whether metabarcoding reads were proportional to eDNA copies, we prepared sets of amplification replicates containing the same amount of an eDNA sample and different amounts of ostrich or emu DNA standard, as illustrated in Fig. 3 (panel 1). Replicates were indexed separately, sequenced, and fish eDNA copies were calculated by comparing fish reads to standard reads in each replicate as follows:

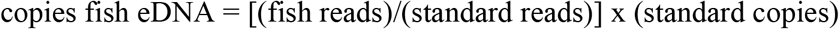

**Fig. 3.**
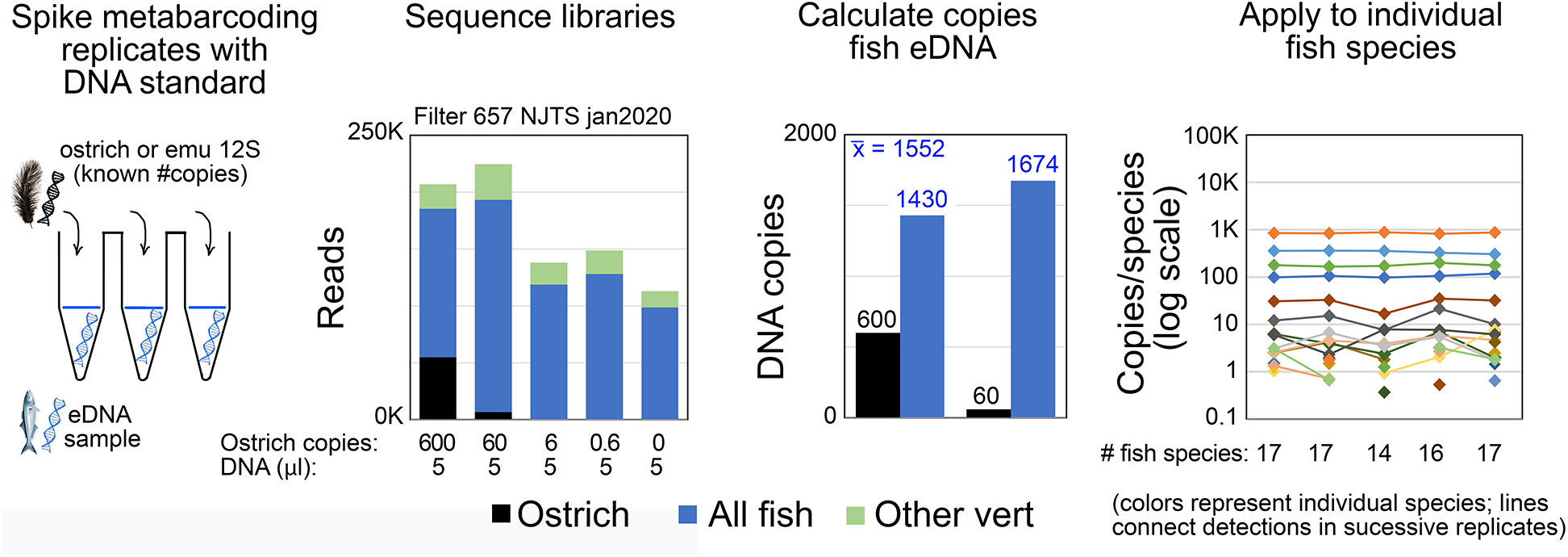
Experimental protocol, representative results. Beginning at left, technical replicates were spiked with known amounts of ostrich or emu 12S DNA. Separate libraries were generated from each replicate, and total fish eDNA copies were calculated by comparing fish and DNA standard reads. At right, eDNA copies for individual fish species were calculated by applying average total fish eDNA copies to each library. Data shown were obtained from one NJOTS sample (source data Supporting Information Tables 4A,4B).

Several findings were evident. First, total vertebrate reads differed among replicates, up to about 2-fold (Fig. 3, panel 2). This might reflect variable efficiency of indexing, PCR inhibition, or other factors. Second, in some replicates prepared with smaller amounts of standard DNA, there were no standard reads (e.g., Fig. 3, panel 2, see replicate with 6 copies ostrich DNA). In these cases, standard reads were present in FASTQ files but were below the filtering threshold (see Methods). Third, replicates containing different amounts of standard DNA yielded similar estimates of fish eDNA copies, consistent with relative reads being proportional to relative DNA copies (Fig. 3, panel 3). Fourth, the calculated copies per species were uniform across replicates for taxa with more than 10 calculated copies (Fig. 3, panel 4). Below that level, some species were detected inconsistently. Inconsistent detection, referred to as pickups or dropouts, is commonly observed in metabarcoding studies (Ficetola et al., 2015). These findings were further explored in experiments reported below, with supportive results.

### Are relative reads proportional to relative DNA copies?

If so, then replicates containing different amounts of standard DNA should yield identical estimates of fish eDNA copies. Replicates containing 5 μl of a DNA sample (n = 18) were spiked with 6000 or 600 copies of ostrich DNA, and copies of fish eDNA in each replicate were calculated by comparing fish reads to standard reads as described above. Different amounts of standard DNA generated similar estimates of fish eDNA copies (Fig. 4, left panel). The average arithmetic deviation from identity was 1.3-fold, and the maximum was 2.0-fold. The direct relationship between relative reads and relative DNA copies held over a wide range of sample eDNA content, from about 50 to about 5,000 calculated copies fish eDNA. This protocol was applied to additional samples (n = 18), except that 600 or 60 copies of ostrich or emu DNA were added. As in prior experiment, different amounts of standard generated nearly identical estimates of fish eDNA copies (Fig.4, center panel). The average and maximum difference in replicate pairs was 1.2- and 1.7-fold, respectively. As above, a proportional relationship between reads and copies was observed in samples with widely varying eDNA content ranging from about 5 to about 5,000 copies fish eDNA. A direct relationship of relative reads to relative DNA copies extended to 60- vs. 6-copy replicates (Fig. 4, right panel).

**Fig. 4.**
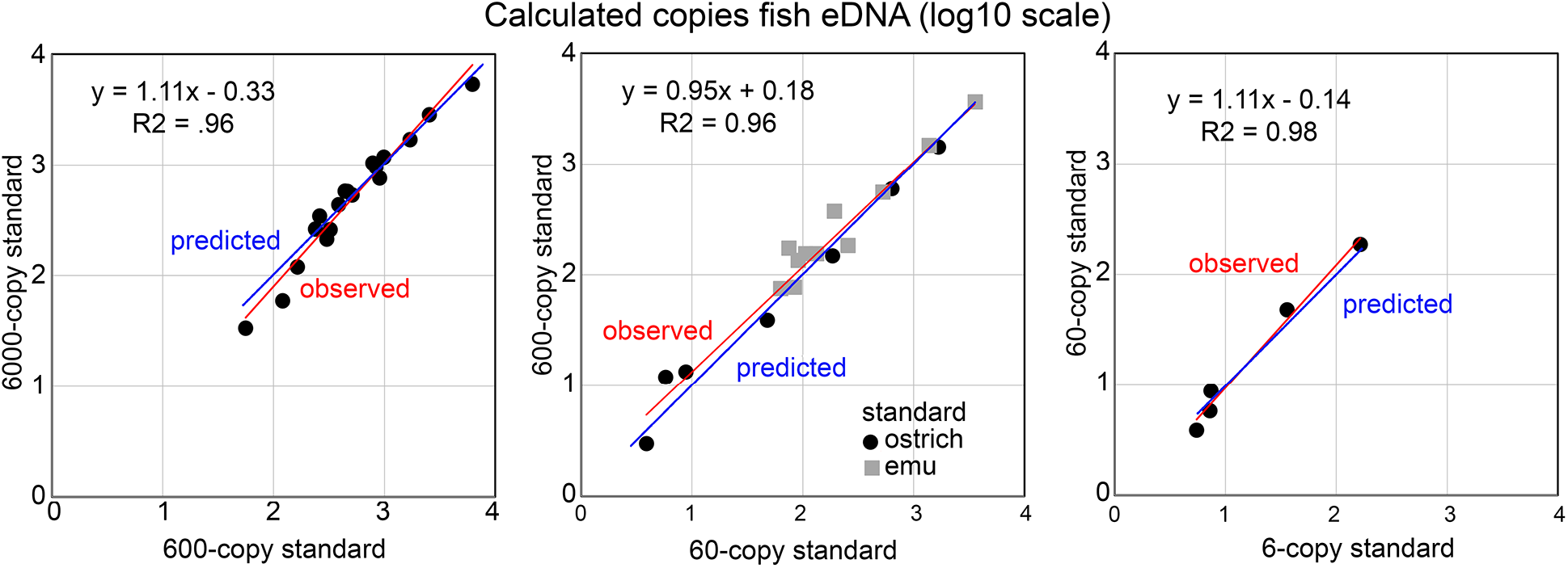
Calculated copies of fish eDNA according to DNA standards. Left: separate libraries were generated from replicates spiked with 6000 or 600 copies of ostrich DNA, and copies of fish eDNA were calculated as described. Each point represents one eDNA sample analyzed with one pair of replicates. Blue line denotes relative reads directly proportional to relative copies. Red line is linear regression of log-transformed experimental data. Center: Protocol applied to a different set of eDNA samples, using 600 or 60 copies of ostrich or emu DNA. Right: Results with subset of samples in center panel, using 60 or 6 copies ostrich standard (source data Supporting Information Tables 4A,5,6).

### Are calculated copies/species proportional to amount eDNA analyzed?

Paired replicates with standards were prepared with 4.375 μl or a 4-fold greater amount, 17.5 μl, of eDNA samples (n = 4). Increased PCR input was reflected in copies/species, but not in reads/species (Fig. 5). Consistent with expectations, there were four-fold more pickups in 17.5 μl replicates, and among all replicates most pickups (21/22, 95%) were eDNAs present at fewer than 10 copies.

**Fig. 5.**
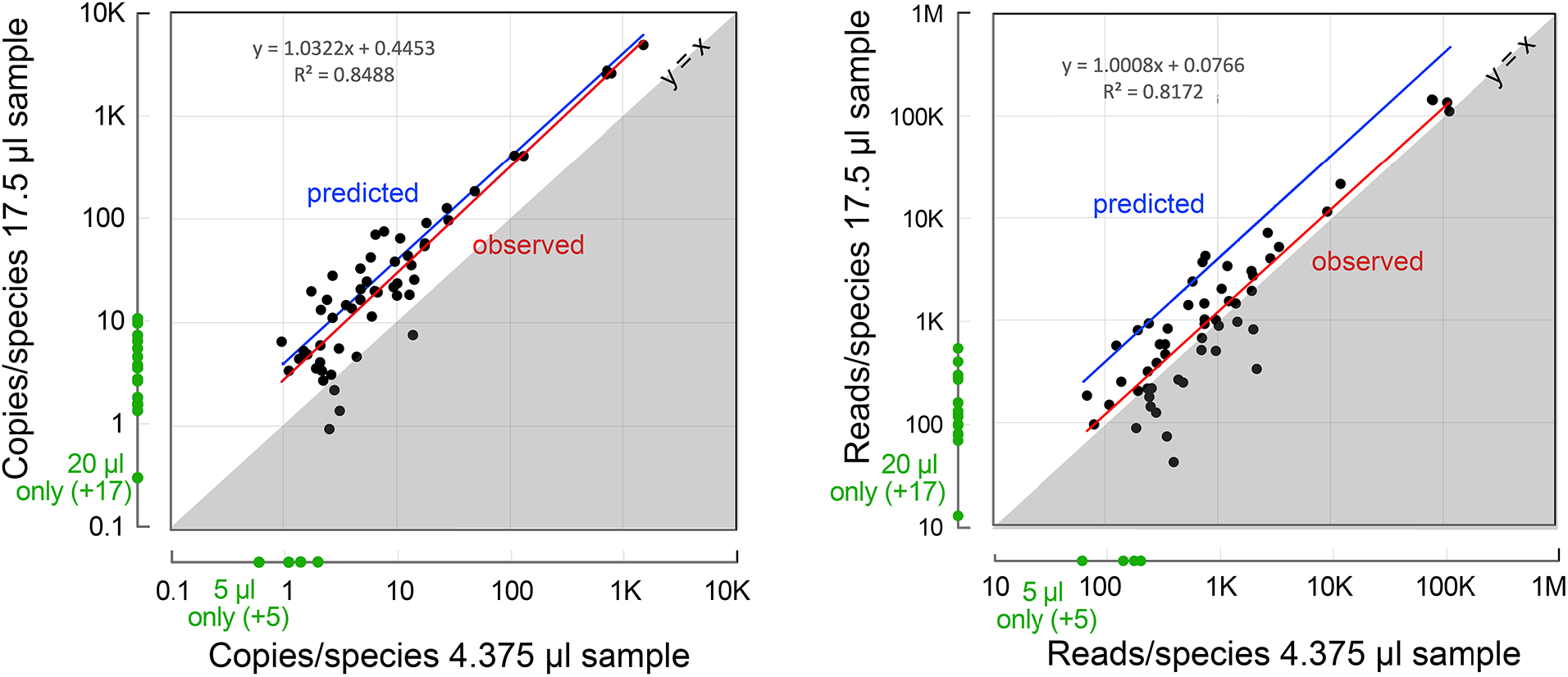
Copies and reads compared to amount eDNA analyzed. Replicates were prepared with indicated amounts of an eDNA sample plus ostrich standards. Each point represents one species in one pair of replicates. Black points represent paired values for copies/species (left) and reads/species (right). Blue line corresponds to copies or reads per species being 4-fold higher in 17.5 μl replicate, red line is linear regression of log-transformed experimental data, and the y = x boundary indicates plot if values were equal. Green points represent pickups, i.e., species detected in one of the paired replicates (source data Supporting Information Table 5).

### Is there a lower limit to reproducible detection?

To further analyze dropouts as illustrated in Figs. 3,5, we compiled detections from samples (n = 8) analyzed with five replicates (Fig. 6). Copies/species and reads/species were calculated by averaging positive values among replicates, i.e., excluding non-detections. Higher eDNA copy number was associated with fewer dropouts (e.g., detected in all replicates: 4^th^ quartile vs. 1^st^ quartile, 97% vs. 0%; p <0.0001, Fisher’s exact test). No dropouts were observed in detections with more than 10 calculated copies (Fig. 6). A similar but weaker trend was observed with raw reads (detected in all replicates, 4^th^ quartile: copies vs. reads, 97% vs. 74%; p = 0.007, Fisher’s exact test).

**Fig. 6.**
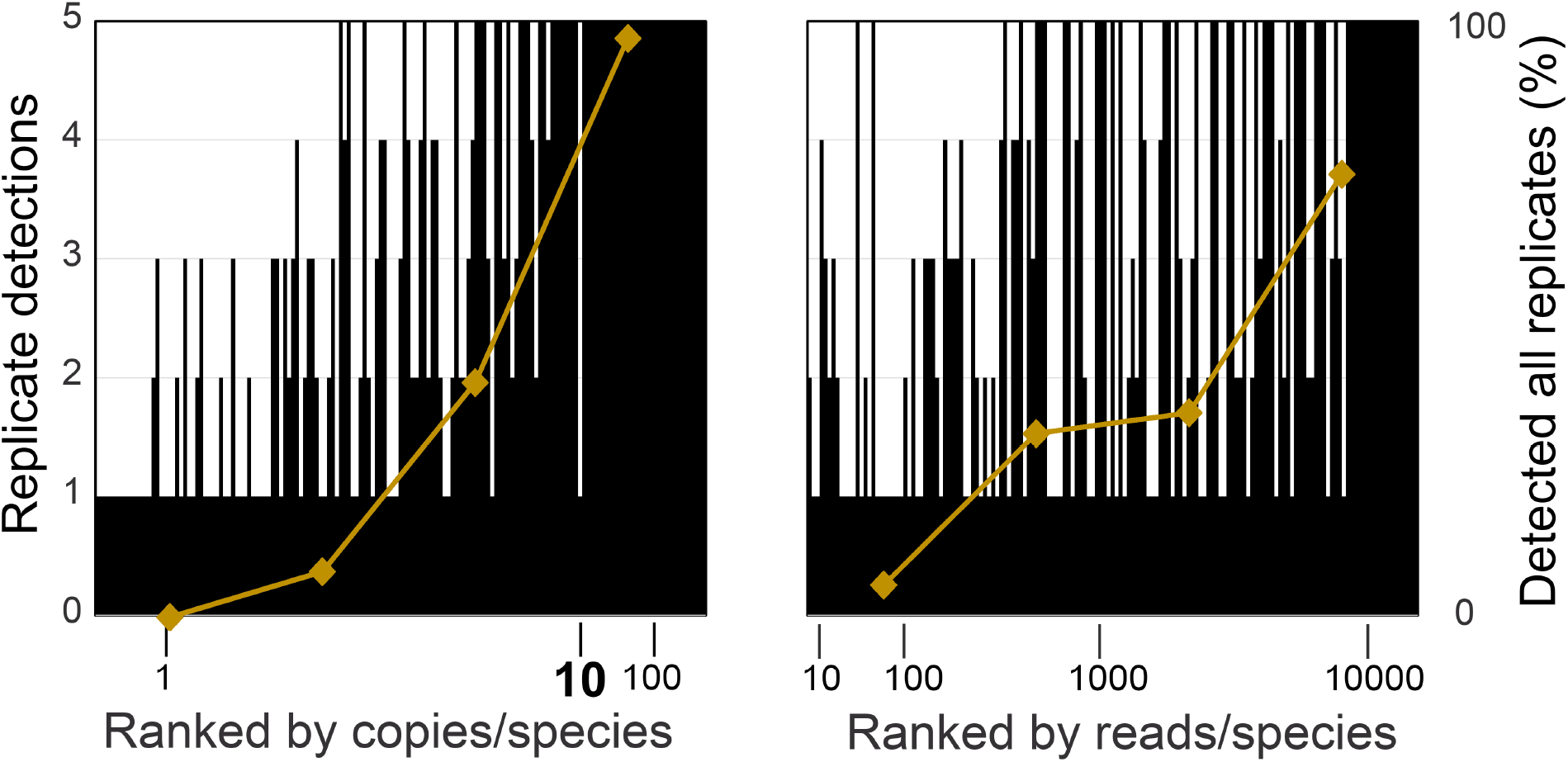
Replicate detection. Each black line represents one species in one eDNA sample analyzed with five technical replicates (n = 154). Superimposed gold line charts percent detections present in all replicates for each quartile (source data Supporting Information Table 4C).

### Does standard correct for distortion of read counts due to non-fish vertebrate DNA?

We analyzed copies in replicates with very different proportions of fish vs. non-fish vertebrate reads, the latter including those due to DNA standard itself (Fig. 7). Despite these differences, fish copies and fish species detection were consistent within replicate sets. Calculated fish copies were higher in replicates prepared from a larger volume water sample, as expected, whereas reads were not (Fig. 7, columns 2,3). Most non-fish vertebrate DNA reads not from standards were human or domestic animal, and the remainder were derived from non-fish wildlife (Supporting Information Tables 4A,5,6).

**Fig. 7.**
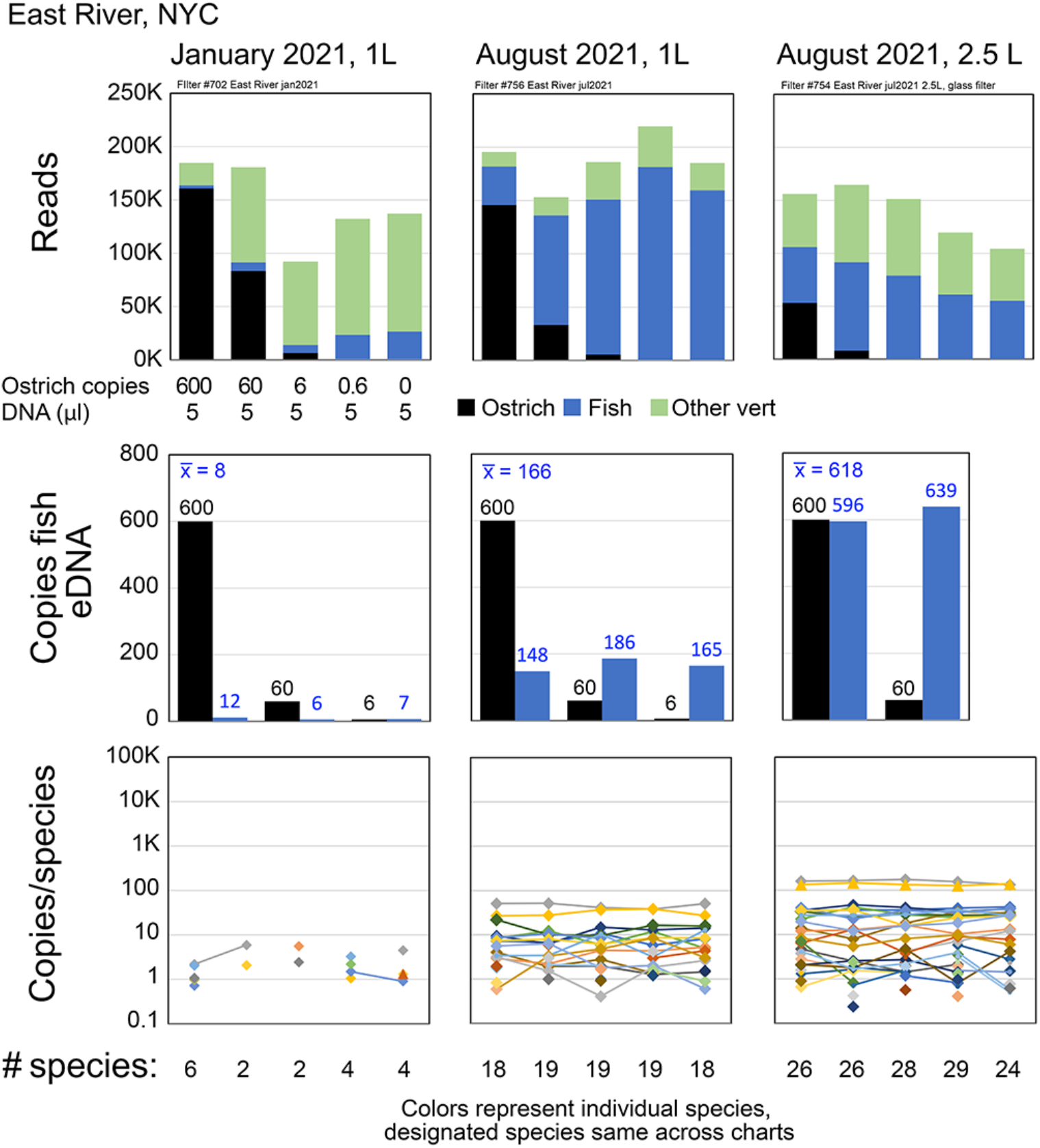
Evaluating distortion due to non-fish vertebrate reads. eDNA samples were analyzed with protocol illustrated in Fig. 3. See text for discussion (source data Supporting Information Tables 4A,4B).

### How important is PCR bias?

We reasoned that if primer bias or PCR efficiency differed among species, then primer sets targeting different gene regions should give different results. Conversely, similar results with different primer sets would be evidence of relative absence of primer or PCR bias, at least for the species present in the sample. To test this hypothesis, we compared results obtained with Riaz primers to those generated with modified MiFish primers (see Methods), which amplify an adjacent region of vertebrate 12S gene (Fig. 2). Libraries were generated from NJOTS samples (n = 19) from January 2020, and the results were pooled. Reads/species were well-correlated between the two primer sets (Fig. 8), although there were greater differences below about 1000 reads/species, which might be due to variability inherent in low-copy number eDNA as noted above (more than 1000 reads, linear regression log-transformed values: slope, 1.00; R^2^, 0.94; fewer than 1000 reads, no significant relationship). The number of pickups was similar between primer sets, and most (10/11; 91%) were represented by fewer than 1000 reads, consistent with these corresponding to low-copy number eDNA.

**Fig. 8.**
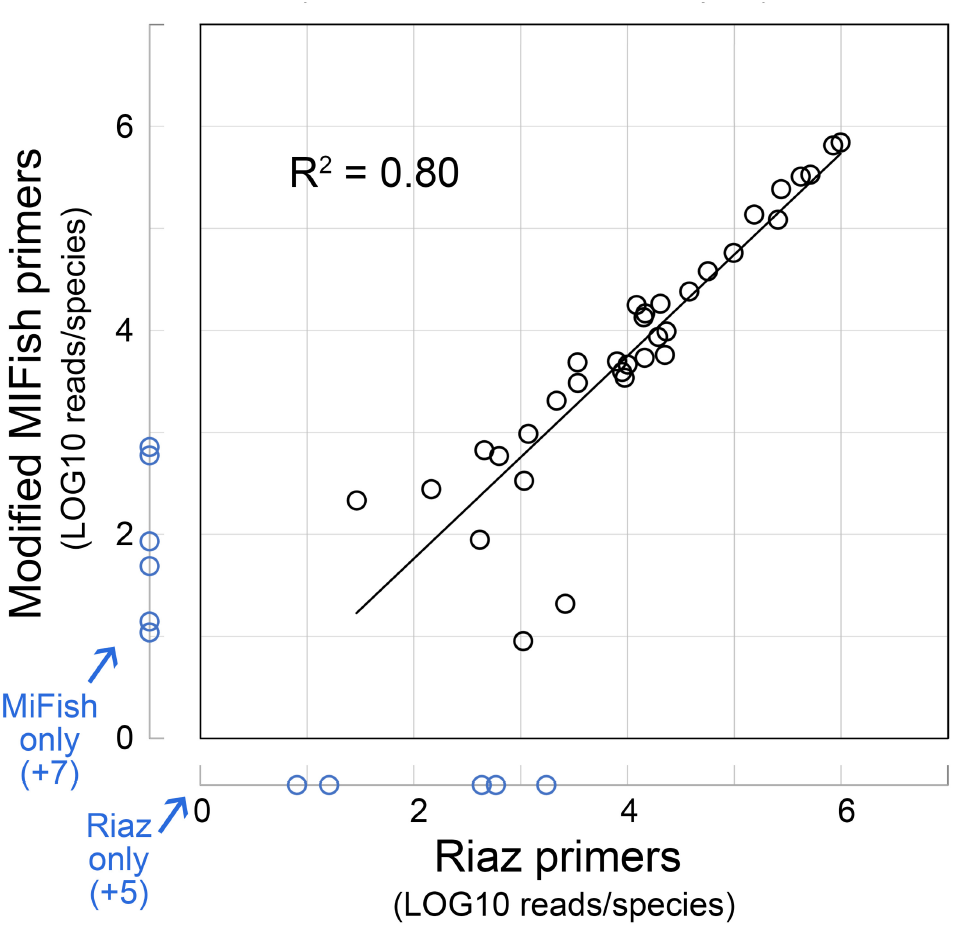
Comparison of reads per bony fish species with different 12S primer sets. Black circles, species with shared detections (n = 33). Blue circles, species detected with one primer set only. Line is linear regression of log-transformed data (source data Supporting Information Tables 7-9).

## DISCUSSION

We tested whether Riaz 12S gene metabarcoding with an internal DNA standard quantifies marine bony fish eDNA. We present several lines of evidence supporting this hypothesis. First, relative reads were directly proportional to relative DNA copies over a 1000-fold range of standard DNA copies (6 to 6,000) and a 1000-fold range of calculated eDNA copies (about 5 to 5,000). Paired replicate measurements differed by an average and maximum of 1.3-fold and 2.0-fold respectively, which was well within our variance target of one order of magnitude. Second, the DNA standard accurately measured the relative proportion of a DNA sample used for PCR, information that cannot be extracted from read data alone. Third, we found a threshold for reproducible detection of about 10 copies/species per PCR reaction and demonstrated improved detection of low-copy number eDNA by analyzing a larger proportion of a DNA sample. These results provide an intuitive mechanistic explanation for the empirical phenomenon of metabarcoding dropouts. Fourth, incorporating a DNA standard corrected for distortion of read counts due to non-fish vertebrate DNA and variation in total reads among replicates. Finally, a comparison of eDNA reads obtained with alternate 12S gene primer sets was consistent with absence of significant differences in amplification efficiency among marine bony fish species.

### Limitations

The bioinformatic pipeline filters out low level detections, those present at less than 1/1000 of total reads per taxon in sequencing run. This threshold aims to reduce effects of tag jumping and sequencing error, However, it may eliminate true positives if total reads for a taxon are high, which was the case for DNA standards. As expected, in some replicates prepared with smaller amounts of standard DNA, there were no standard reads after filtering (e.g., Figs. 3,7), thus we could not use these to test our hypothesis. This limit could be addressed with unique dual indexing (Bohman et al., 2022), which should reduce tag jumping and need for threshold. Although not a factor in this study, on a MiSeq, a high level of DNA standard (>100,000 copies per reaction) suppresses detection of fish eDNA in near shore samples. If desired, this could potentially be overcome with deeper sequencing using NovaSeq or a similar platform (Singer, Fahner, Barnes, McCarthy, & Hajibabaei, 2019).

Comparison of reads/species with MiFish and Riaz primer sets was consistent with relatively modest differences among bony fish species in PCR efficiency. However, we cannot exclude primer/PCR bias, particularly for species not represented in the NJOTS samples analyzed. Both primer sets amplify cartilaginous fish eDNA poorly, and this limitation might apply to other species. It is well-established that broad range metazoan primers can give very different results in terms of taxon detection and relative abundance (Liu & Zhang, 2021; Kumar, Reaume, Farrell, & Gaither, 2022). Metabarcoding mock community samples can enable adjustments that make up for differences in PCR efficiency among taxa (Shelton et al., 2022b). Regarding the latter, it may be of interest to test mock communities that closely mimic eDNA samples, i.e., in which target DNAs are present at eDNA-like concentrations and comprise a small fraction of total DNA. Primer mismatch, which can be evaluated in silico, may be the main factor determining 12S amplification efficiency in marine fish. For example, in Shelton et al., 2022b, in analyzing fish eDNA with MiFish-U F/R primers, outliers were cartilaginous species, which are known to amplify poorly with this primer set, while teleost taxa amplification was relatively uniform. Recent reports demonstrate good correlation between relative reads and relative organism abundance, which implies that even without internal standards, metabarcoding correctly reports relative eDNA abundance in some settings (Afzali et al., 2020; Di Muri et al., 2022; Ershova, Wangensteen, Descoteaux, Barth-Jensen, & Præbel, 2021; Klymus, Marshall, & Stepien, 2017) The reverse MiFish primer overlaps with the forward Riaz primer, which might mean there is less difference in amplification efficiency between these primer sets than one would otherwise expect. PCR bias could be further evaluated by testing both primer sets on samples containing eDNA of other species and on mock communities, calibrating modified MiFish primers with protocols similar to those here, and comparing NGS copies to results with single-species qPCR or ddPCR assays.

Copy number calculations in this study assumed that the ostrich and emu amplicons employed as DNA standards were accurately measured and that they amplified with the same efficiency as fish eDNA. Regarding the former, dilution experiments analyzed with Poisson distribution indicate these standards are within about 3-fold of nominal value (Stoeckle et al., 2022). Similar support is provided by observation that the lowest values for calculated copies/species per replicate were regularly at or close to 1 (e.g., Figs. 3,6,7). The latter aspect, relative amplification efficiency, could be evaluated by adding standards to mock eDNA community samples.

### Other reports

A related approach using internal standards to quantify MiSeq metabarcoding of marine fish eDNA is described by Ushio et al., 2018. In that report, eDNA samples were spiked with four internal standards present at 50 to 1000 copies, amplified in triplicate with unmodified MiFish-U primers, and resulting amplicons were pooled, indexed, and sequenced on a MiSeq. eDNA copies were calculated by comparing fish reads to standard reads, and these values were compared to qPCR copies for total fish eDNA and two fish species. Some samples amplified poorly or not at all, possibly due to PCR inhibition, and correlations with qPCR were modest even after excluding outliers. The authors note that off-target amplification with unmodified MiFish-U primers may have interfered with qPCR assay for total fish eDNA. In addition, we observe that calculated fish eDNA copies/species were mostly below 10 copies per reaction, which may have hampered reproducible measurement. This approach has been applied to quantify marine fish eDNA around artificial reefs (Sato et al., 2021), but to our knowledge accuracy has not been further evaluated.

### Looking ahead

Our findings support further testing of the methodology outlined here for marine bony fish metabarcoding, namely, incorporation of an internal DNA standard and separate analysis of replicates. A reasonable starting protocol would be a pair of replicates for each eDNA sample, one with 600 and one with 60 copies of DNA standard. Potential avenues for further investigation include comparing metabarcoding eDNA measurement to that obtained with qPCR, applying MiFish and Riaz primers to additional samples, testing different DNA standards, and adapting this approach to analyze cartilaginous fish eDNA. In the long run, just as a net catches some fish while others swim away, a metabarcoding assay does not need to perfectly report eDNA abundance to be useful, but it does need to be reproducible over a range of eDNA concentrations. Incorporating a DNA standard will likely help. In evaluating how eDNA abundance relates to fish abundance, it may be helpful to consider allometric scaling (Stoeckle et al., 2021, Yates et al., 2020).

Our results add evidence that current metabarcoding protocols suffice for relatively abundant marine fish eDNA, which presumably includes that of most commercial species. Conversely, we find that eDNA rarity is the biggest challenge to fish eDNA metabarcoding. With a typical protocol applied to near shore mid-Atlantic samples, eDNA of many bony fish species was below the limit of reproducible detection. This could be addressed by amplifying a larger proportion of a DNA sample, starting with multiple or larger water samples, or utilizing a different collection strategy such as targeted collection near to organisms, passive collection, or examining DNA trapped by filter feeders (Baker et al., 2018; Bessy et al., 2020, 2021; Hunter, Ferrante, Meigs-Friend, & Ulmer, 2019; Mächler, Deiner, Spahn, & Altermatt, 2016).

### Summary

Effective ocean management relies on accurate, up-to-date information on marine life. eDNA provides a relatively low-cost, harmless, widely applicable, and potentially autonomous method that will supplement traditional ocean survey techniques. Reliable measurement of eDNA concentration will enhance the value of this new technology.

## SUPPORTING INFORMATION

Supporting Information Tables 1-9

Table 1. NJ Ocean Trawl Survey, monthly totals, Jan2020

Table 2. Primers, PCR parameters

Table 3. Match Riaz, MiFish IDs

Table 4A. Ostrich 1A: 8 samples, 5 replicates

Table 4B. Ostrich 1B: copies per species

Table 4C. Ostrich 1C: compare detections vs reads, copies

Table 5. 4.375 vs 17.5; Emu 600, 60

Table 6. Ostrich 6000, 600

Table 7. Riaz NJOTS jan2020

Table 8. MiFish NJOTS jan2020

Table 9. MiFish vs Riaz, NJOTS jan2020

Supporting Information File 1. FASTA file Riaz 12S gene reference sequences

## DATA ARCHIVING

Illumina FASTQ files underlying this article are deposited in NCBI Bioproject ID PRJNA793893.

## ACKNOWLEDGEMENTS

This work was supported by NOAA Grant NA20OAR01104XX. Sampling during the New Jersey Ocean Trawl Survey was supported by US Fish and Wildlife Service Wildlife and Sport Fish Restoration Program. We thank Greg Hinks and Stacy VanMorter at Bureau of Marine Fisheries, New Jersey Department of Environmental Protection for sharing NJOTS catch data. At The Rockefeller University we thank Jeanne Garbarino, Disan Davis, and Odaelys Walwyn for laboratory resources and assistance. The funders had no role in the study design, data collection and analysis, decision to publish, or preparation of the manuscript.

